# Cleavage region organizes the structural architecture of the B2 SINE ribozyme

**DOI:** 10.1101/2024.07.22.604665

**Authors:** Ankush Singhal, Tyler Mrozowich, Susmit Narayan Chaudhury, Carlos Rivera, Maulik Badmalia, Jeannie T. Lee, Trushar R. Patel, Karissa Y. Sanbonmatsu

## Abstract

The SINE-encoded B2 retrotransposon is an RNA Polymerase III transcript that gets upregulated during various cellular stress responses. The B2 noncoding RNA can directly inhibit RNA Polymerase II, leading to a significant downregulation of transcripts during stress. Our recent findings have shown that B2 is a self-cleaving epigenetic ribozyme and that cleavage can be induced by interactions with epigenetic factors, co-regulating its function across distinct chromatin-binding target loci. Here, by integrating RNA chemical probing, small angle X-ray scattering, and 3D motif modeling, we determine structural ensemble-to-function relations for the B2 SINE ribozyme RNA. Perturbations of the RNA suggest that the B2 SINE ribozyme has a well-defined secondary structure and dynamic tertiary structure that both critically depend on the presence of the active site of cleavage. In an RNA engineering approach, we examine the effect of point mutations, deletions of the main cleavage site and deletions of the cleavage domain on the structural ensemble of the RNA. By combining this with functional data, we obtain relations of structural ensembles to various functional states. This perturbative approach serves as a template to unravel the relation of structural ensembles to functional states for other ncRNA and mRNA systems.

## Introduction

Complex Eukaryotic genomes contain tens of thousands of repetitive sequences originating from transposition reactions. Once considered junk fragments or parasites of the genome, we now know transposable elements (TEs) are critical regulators of many biological phenomena^1, 2^. The major class of TEs, retrotransposons, mobilize their DNA through an RNA intermediate, causing duplication of the target sequence. Short interspersed nuclear elements (SINEs) account for half of the repetitive sequences in humans and mice. Despite coming from a non-autonomous origin, they require trans-acting enzymatic machinery for achieving transposition, provided by proteins encoded in long interspersed nuclear elements (LINEs).

Together with primate Alu elements, the murine B2 repeats were the first SINEs to be cloned and sequenced about four decades ago^3, 4^. They are the most studied class of SINEs in mice, represented in more than >120.000 copies in their genomes. RNA polymerase III (RNA pol III) and RNA Polymerase II (RNA Pol II) transcribe B2, 180 nucleotides long RNAs as internal sequences of longer RNAs^5^. Interestingly, B2 RNA is actively transcribed in embryonic stem cells and during the early stages of embryogenesis, with B2 levels declining as development progresses towards somatic tissues^5, 6^ Transcription changes are affected by epigenetic silencing mechanisms that inhibit RNA Pol III–mediated transcription, such as DNA CpG methylation^7^, positioning of linker histones^8^, and histone H3 lysine 9 trimethylation (H3K9me3)^9^ Remarkably, the transcription of B2 elements gets upregulated upon different cellular stressors such as heat shock^10, 11^, translational inhibition^12^, viral infection^13, 14^, genotoxic agents^15^, ethanol, and others. A recent report also showed the upregulation of B2 RNAs in mouse amyloid beta pathologies^16^. These reports underscore B2 RNA as central to the mammalian stress response.

B2 SINEs contain an RNA Pol II promoter that can transpose to new regions, establishing novel transcripts, such as Lama3^17^, and B2 RNAs are identified as eventual up-regulators of translation^18^. B2 also harbors CTCF binding motifs, a vital feature in regulating the establishment of chromatin boundaries between silent and active territories^19, 20^. In addition to these molecular functions, breakthrough discoveries in the early 2000s showed that B2 RNA is required explicitly for downregulating RNA Pol II-dependent genes during heat shock^21^. Mechanistically, B2 can bind directly and reversibly to RNA polymerase II with nanomolar affinity, substantially inhibiting its transcriptional activity and preventing it from forming initiation complexes^22^. Furthermore, B2 restricts the physical interaction between RNA Pol II and the DNA regions at the core promoter of the substrate^23^. The binding and inhibitory functions were ascribed initially to a domain located within the 81-131 nt region, with further analyses showing the minimal region capable of binding – but not inhibiting – RNA Pol II is within 99-131 nt, whereas the 105-108 nt segment exerts transcriptional repression^24^. Additionally, B2 harbors an Mg^2+^-dependent self-cleaving ribozyme activity between positions 81-124, whereas nucleotides 100-101 are determinants of the self-cleaving pattern specificity^25^. Thus, B2 is categorized as a SINE-encoded self-cleaving ribozyme that plays a specific regulatory function in transcription both in resting conditions and in response to cellular stress, working together with epigenetic modifiers to regulate different steps of Pol-II-mediated gene expression.

Despite its biological significance, this versatile RNA’s three-dimensional structure remains largely unknown. An initial assumption was that ncRNAs are flexible and can have dynamically changing conformations^26, 27^. Interestingly, it has been shown that ncRNAs can adopt modular secondary structures by chemical probing techniques. Selective 2’-hydroxyl acylation analyzed by primer extension (SHAPE)^28^ evaluates the local backbone flexibility of RNA molecules at single-nucleotide resolution in diverse solution conditions, and there is evidence that ncRNAs preserve functionally relevant sequences and structure and form unique interacting domains^29, 30^. However, an RNA’s functionality relies not only on primary or secondary sequence but also its tertiary conformation. The one-dimensional (1D) and three-dimensional (3D) data extracted from small-angle X-ray scattering (SAXS) provide global structural information about molecules in the solution. Therefore, SAXS is an excellent complementary approach to solving RNA structure synergistically with other biophysical methods. Numerous studies have used SAXS to help determine several RNA^31–33^ and RNA-protein complexes structures^34^.

Here, we present an interdisciplinary approach that uses native purification, chemical mapping, SAXS, and computational modeling to elucidate the sequence-to-structural ensemble relationship of B2 RNA. Comparing the wild-type B2 RNA to variants of B2 with reduced catalytic activity, we find significant changes in both secondary and tertiary structures. We find the wild-type RNA to have a well-defined secondary structure but a flexible and dynamic tertiary structure. When deleting the cleavage site, known to eliminate the activity of the RNA, the secondary structure is reorganized while the tertiary structure becomes less mobile. When we delete the entire cleavage domain, the 5’-domain remains intact, while the 3’ domain may undergo oligomerization with other B2 SINE RNAs. Point mutations that allow cleavage but weaken activity substantially leave the secondary structure intact but likely rearrange tertiary contacts. These insights provide a structural basis for understanding the biological properties of B2 RNA and shed light on its functional implications.

## Results and Discussion

Our work reveals a stable, denaturant-resistant structure that contains domains, highlighting the presence of structured and unstructured regions, including six stem loops and an A-rich 3’ end with enormous flexibility, which is noticeably different when comparing the consensus WT B2 structure with a B2J, a natural, two nucleotide mutant variant. B2J shows weak self-cleaving activity^35^ which shows different structural features with RNA sequences. B2J differs from WT by (i) an A to C change at position 55 and (ii) a U to C change at position 114. Two distinct deletion mutants of the cleaving region, with targeted deletion of active sites, were also examined, B2Δ96-105 and B2Δ81-124.

### Transcription and Purification of B2 RNA

The B2 RNA variant *mm1a* exhibits both transcriptional inhibition properties and self-cleavage activity^35, 36^. We transcribed it via T7 RNA polymerase from a PCR product containing a T7 promoter at the 5’ end and purified it via size-exclusion chromatography under native conditions, not including any organic solvent that could affect the post-transcriptional conformations of B2 (Figure 1A). SEC purification of B2 RNA produced a large primary peak enriched in the full-length B2 RNA and a secondary peak containing partially cleaved B2 species, as seen in denaturing Urea PAGE (Figure 1b, 1c), suggesting the ribozyme acquires its active conformation as soon as it is being transcribed. We further confirmed the activity of the B2 ribozyme, as seen in a time-course experiment performed in the presence of Mg^2+^ as a canonical inorganic cofactor (Figure 1d). Intriguingly, B2 self-cleavage activity resisted multiple nucleic acid denaturants, such as betaine, formamide, and urea, suggesting a high structural stability of the active conformers of this ribozyme (Figure 1e). After validating the compatibility of this RNA native purification methodology with ribozyme activity assays, we employed it to purify B2J, a naturally occurring B2 variant whose activity follows a different cleavage pattern compared to the wild-type B2 used in this study (Figure 1f). In addition, we purified two mutant versions of B2, lacking the leading site of cleavage that is detectable *in vivo* (B2(Δ96-105)) and lacking a critical region for its catalytic activity (B2(81-124) (Figure 1f))^35^.

**Figure 1:**
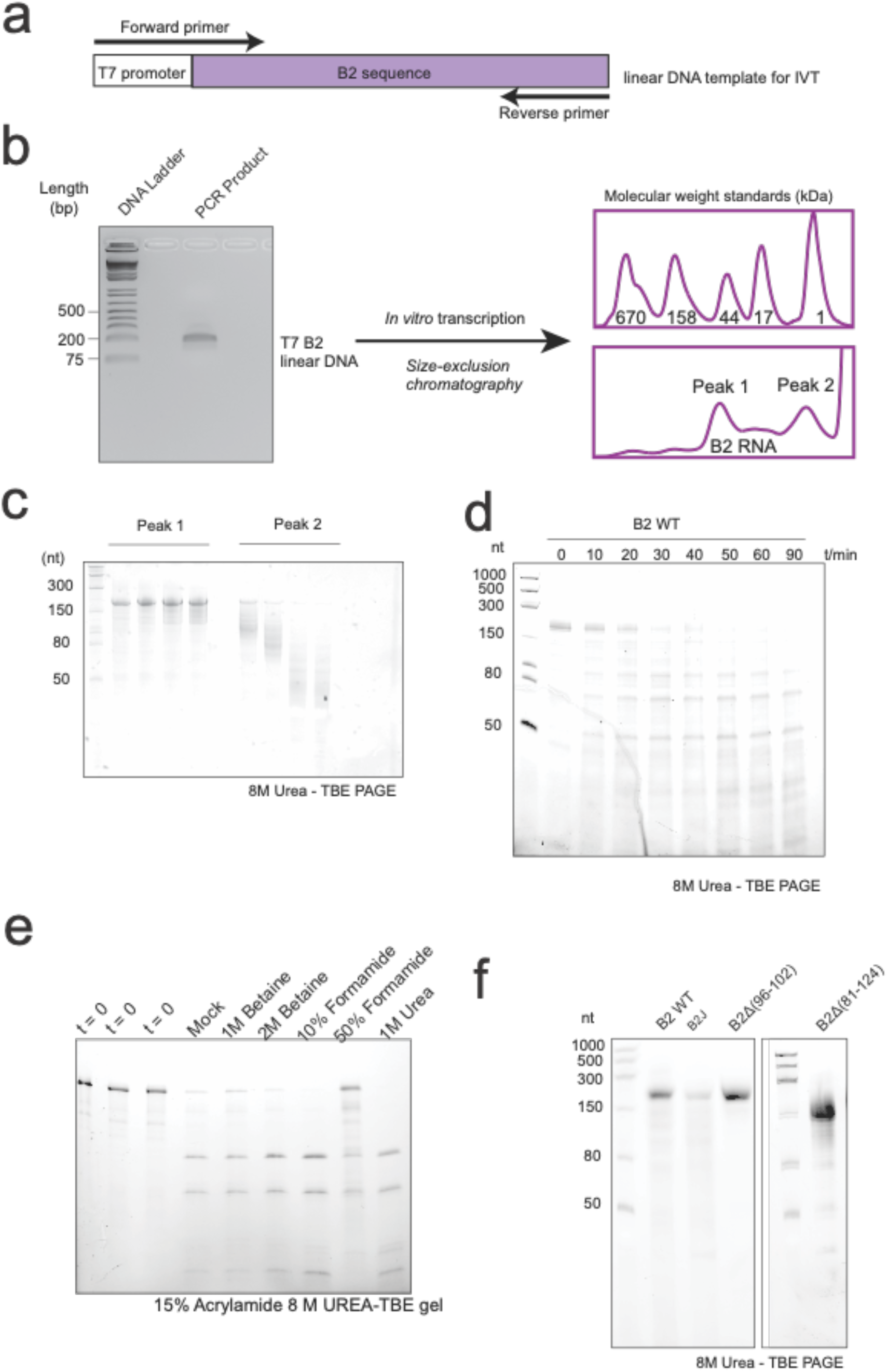
The B2 self-cleaving activity resists denaturing conditions. A and B panels depict the strategy to synthesize the B2 ribozyme by *in vitro* transcription and size exclusion FPLC. (C) B2 size exclusion chromatogram fractions depicting the standard cleavage pattern. (D) Time course of B2 self-cleaving activity.(E) B2 self-cleavage activity against multiple nucleic acid denaturants (F) B2 mutant constructs compared to WT under denaturing conditions.

### Secondary structure determination via chemical probing

Previous work has shown that nucleotides 3 to 149 are the functional portion of the 178 nt B2 RNA, in which regions 3 to 74 nts are known not to bind to polymerases nor repress transcription^22^. Detailed structural information is needed to gain further functional insight into the mechanism of B2 RNA. We used chemical probing to perform a structural deletion analysis of the B2 RNA to deduce the secondary structure for B2 RNA using our SHAPE probing data. SHAPE electropherogram experiments of B2 RNA and its mutant variants were performed in the presence of 0.5 mM Mg^2+^ and at 25°C. The SHAPE electropherogram for WT-B2-RNA (Figure 2A) represents three distinctive structural domains, ranging from 1-72 nts, 73-153 nts, and 154-178 nts.

**Figure 2:**
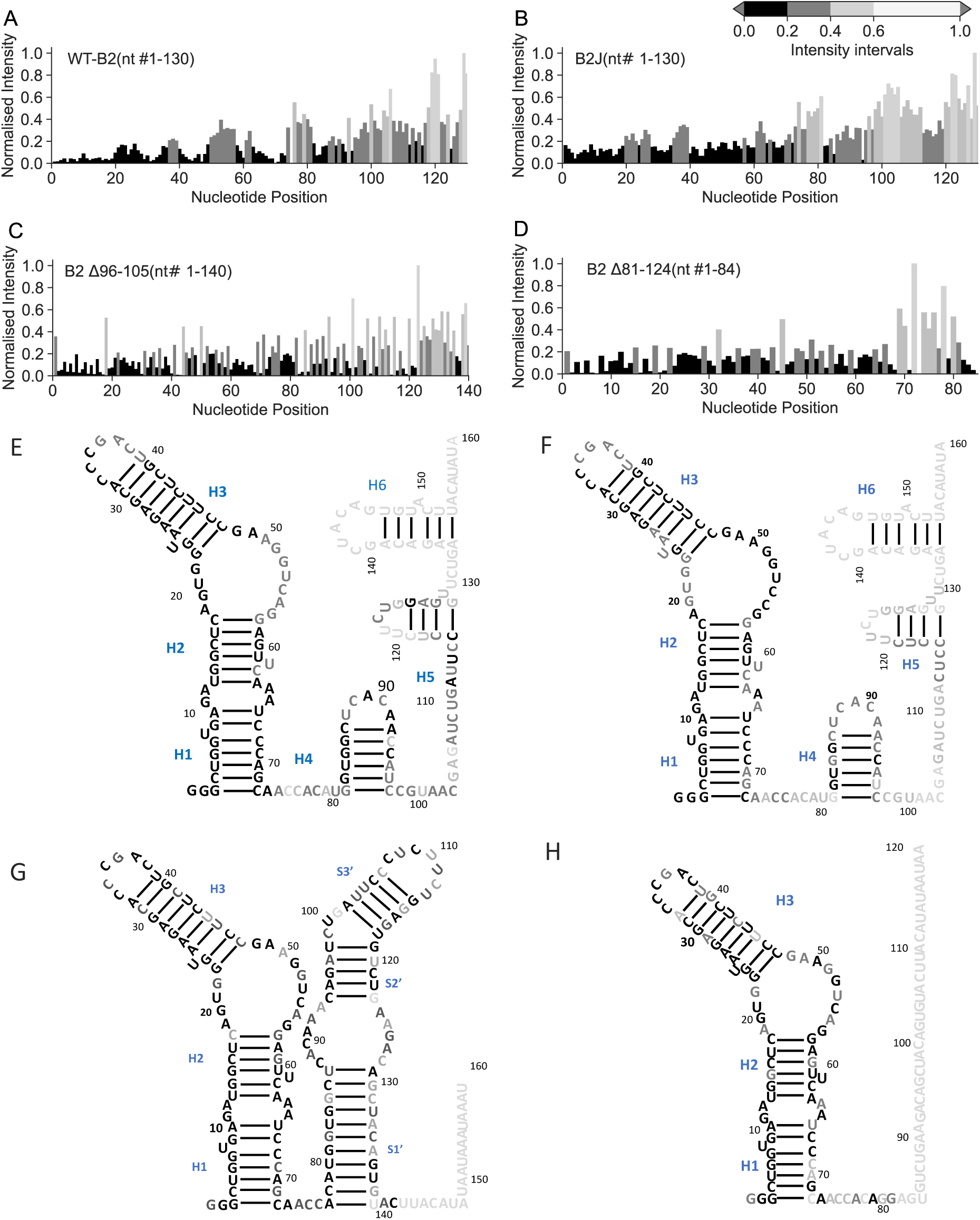
Chemical probing data reveals the secondary structure of B2 and its variant (A) SHAPE electropherogram of WT B2 RNA is performed in the presence of 0.5 mM Mg^2+^ ion. The vertical segments indicate the three structural regions of the RNA. Nt#1-72 is the most structured region of RNA, as indicated by very low SHAPE reactivities. (B) SHAPE electropherogram of B2J RNA. (C) SHAPE electropherogram of 178 nucleotides B2Δ96-105 RNA. (D) SHAPE electropherogram of 178 nucleotides B2Δ81-124 RNA. The electropherogram corresponds to the SHAPE probing experiment of truncated RNA sequences for all the cases. The low SHAPE reactive regions indicate the presence of structured regions, whereas nucleotides with high SHAPE reactivities indicate interhelical junctions. Based on our SHAPE data, we have deduced the secondary structure of (E) WT B2, (F) B2J, (G) B2 Δ96-105, and (H) B2Δ81-124. The unstructured regions are not shown for clarity.

Domain I from 1-72 nts forms a long arm consisting of three stems, two bulges, and an end loop (Figure 2B). Following the nt #72, we notice increased structural flexibility, as suggested by enhanced peak intensities. These enhanced peak intensities indicate the existence of a disordered region. Following the region, a brief reduction in the structural flexibility of nucleotides occurs, which supports the presence of a standalone helix (H4). A relatively long inter-helical junction is also observed immediately after the helix. We observe two more helices (H5 and H6) and a short stretch of inter-helical junctions. The second domain (nt# 73–153) consists of three helices (H4-H6) and three inter-helical junctions, see Figure 2B. At the 3’ end of the WT B2 RNA, we observe an abrupt increase in overall SHAPE reactivity, suggesting an extended, highly disordered region (see Supplementary Figure 1A). Domain III from 154-178 nts is a 25 nts long single-stranded tail. The deduced secondary structure aligns with the previously published results^24^.

The point mutated B2 WT variant, B2J, shows an almost identical secondary structure to B2 WT. B2J contains a two-nucleotide mutation, a change at nt #55 (A to C) in the stable domain 1, and a 2nd mutation at position 114 (U to C) in the inter-helical region (Figure 2C). These point mutations cause no drastic changes that can be observed in B2J’s SHAPE electropherogram (Figure 2B) and result in no change to the secondary structure compared to wild-type. Similar to WT B2 RNA, we observe an abrupt increase in overall SHAPE reactivity at the 3’ end of the B2J RNA, suggesting an extended, highly disordered region (see Supplementary Figure 1B). This finding was surprising, as B2J contains a noticeable reduction in B2 cleavage, which leads us to believe that this biochemical difference might be due to a change in tertiary architecture, which might not be visible through chemical probing. Previous research has indicated that a specific portion of domain 2, spanning nucleotides 75 to 149, is likely sufficient to act as a transcriptional repressor^24^ Additionally, we previously showed that deletion mutations to this domain cause changes to the B2 self-cleavage^35^, and therefore, we sought to investigate the structural implications of these mutations (see Supplementary Figure S2). Chemical probing experiments were conducted identically to B2 and B2J, and the resulting electropherograms and secondary structures are presented in Figure 2 E and Figure 2 F, respectively. We observe a notable divergence in the secondary structures for both the deletion mutants compared to WT.

The secondary structure of domain 1 (nucleotide positions 1 to 72) remained relatively consistent in both mutants, regardless of the deletions, and matches the original B2 WT, containing regions H1, H2, and H3 (Figures 2 E-F). However, the B2 deletion variant, B2Δ96-105, which has reduced catalytic activity^35^, displays the emergence of a stable elongated arm spanning position(s) 76-140. This arm has three stems (designated S1’-S3’) and three bulges (Figure 2 G). Interestingly, this deletion occurs primarily in a single-stranded stretch of B2 and profoundly affects the secondary structure. Notably, B2Δ96-105 still includes a 3’ disordered region spanning 20 nucleotides (see Supplementary Figure 1C), similar to B2 WT and B2J. Previous mutagenesis experiments show that nucleotide region(s) 45-55 and 100-101 influence B2 activity in the presence of EZH2^25^. Surprisingly, these regions display high secondary structural conservation across all B2 mutants. The deletion of the cleavage domain resulting in the B2Δ81-124 mutant, which has no detectable cleavage activity, led to another significant alteration in the secondary structure of the RNA molecule. Unlike B2Δ96-105, which restructured the nucleotides following nt # 1-72 into an elongated arm, the SHAPE electropherogram for B2Δ81-124 indicates an extensive unstructured region following domain 1 (Figure 2 H and Supplementary Figure 1D). B2Δ81-124 does not contain any other notable secondary structural features.

### Small angle X-ray scattering structural studies at low-resolution

Following chemical probing, we needed to assess if the overall tertiary structure was consistent throughout the B2 variants. Evaluating the structural complexity of each RNA, we believe that the inherently disordered regions evident in all B2 variants would prove incompatible with high-resolution techniques like X-ray crystallography or CryoEM. Therefore, we assess the solution structure for each B2 variant in low resolution via SAXS. SAXS was performed on the four different variants of B2 RNA to surmise if the mutations or deletions would impact the overall tertiary architecture of the RNA. Raw scattering data is represented as relative intensity vs. scattering angle (see Figure 3A). Raw scattering data was then transformed and subjected to Guiner analysis (see Figure 3B). Linear regression of each Guiner analysis indicated that all four collected data sets are free of electrostatic interactions, which could affect molecular interactions. B2 RNA has a radius of gyration of 56.5 Å, consistent with an expected Rg of a folded RNA of 178 nucleotides in length (see Table 1). Expectedly, B2J has a similar Rg value of 55.6 Å, likely because the difference between B2 and B2J RNA is a two-nucleotide mutant variant. These two results were consistent with our chemical probing assays,which indicates that B2 and B2J do not show a noticeable difference in secondary structure. When comparing the Rg of the deletion variant B2Δ96-105, we also do not observe a significant change, which can be attributed to the deletion being only nine nucleotides. The most prominent difference is when comparing B2 to the second deletion mutant, B2Δ81-124. B2Δ81-124 results in a drastic change in Rg, an increase to 78.21 Å. This increase (instead of a decrease) is likely due to the dimerization of the B2Δ81-124 RNA, which could result from the increasing disorder of the secondary structure, evident in SHAPE analysis (Figure 2E).

**Figure 3:**
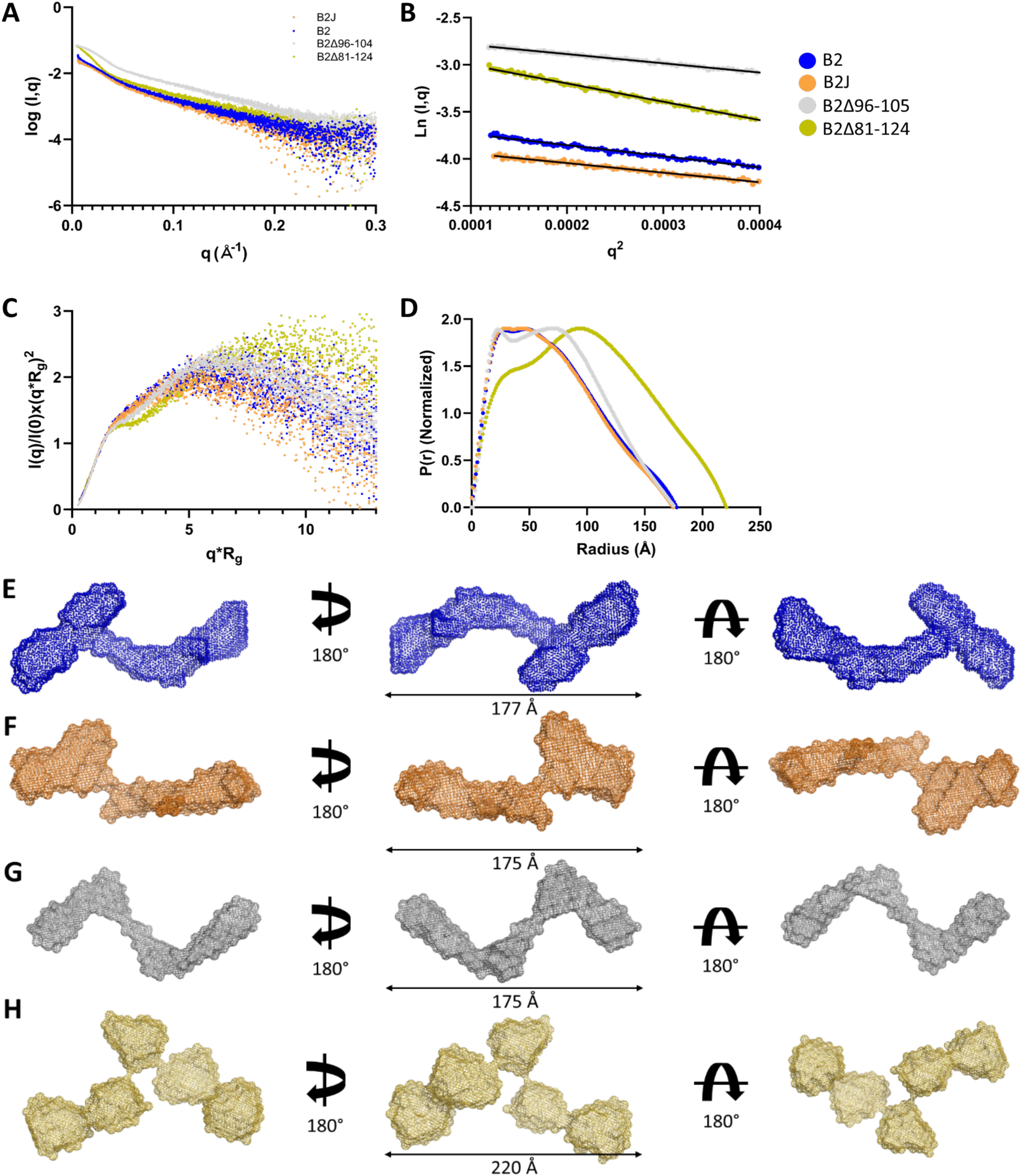
Evaluation of B2, B2 mutations, and truncated RNA through SAXS (A) scattering intensity (Log I(q)) versus scattering angle (q = 4π sinθ/λ) of merged SAXS data for all four RNA. (B) Guinier plots (plot of ln(I(q)) versus q^2^), which allow for the determination of Rg from the low-angle region data. (C) Dimensionless Kratky plots (I(q)/I(0)*(q*Rg)^2^vs q*Rg) for all four RNA samples demonstrating relative foldedness of the extended structures. (D) Pair-distance distribution (P(r)) of all four RNA samples representing D_max_ (maximum particle dimension) and real-space Rg. Ab initio reconstructed three-dimensional models for B2 RNA. The filtered average model is shown and rotated 180 degrees in the X and Y coordinates. Solutions structures of various RNAs are depicted in (E) B2 RNA, (F) B2J RNA, (G) B2Δ96-105, (H) B2Δ81-124.

**Table 1:** Structural characteristics of B2 and its three variants. D_Max_ is the radius when the P(r) distribution approaches zero. Rg is the radius of gyration calculated using the P(r) function. The χ ^2^ (SAXS) value determines the correlation between the experimental and model-derived SAXS data. The χ ^2^ (Model) value determines the goodness of the fit between the averaged SAXS and over the top 5 all-atom models.

| Name | $D_{\max}$ | $\chi^2$ (SAXS) | $\chi^2$ (Model) | $R_g$ (SAXS) | $R_g$ (Model) |
| --- | --- | --- | --- | --- | --- |
| WT-B2 | 177 Å | 1.60 | 3.69 | 56.25 Å | 52.33 Å |
| B2J | 175 Å | 1.39 | 3.25 | 55.60 Å | 54.23 Å |
| B2Δ96-105 | 175 Å | 1.11 | 6.93 | 56.68 Å | 52.90 Å |
| B2Δ81-124 | 220 Å | 1.12 | 6.78 | 78.21 Å | 72.61 Å |

We performed dimensionless Kratky analysis to evaluate the relative foldedness of each RNA. An unfolded molecule will display a sharp initial increase, followed by data that continues upward. Three RNA species (B2, B2J, and BΔ96-105) show a plot that trends towards returning to zero (Figure 3C), indicating the RNA are folded solution, but likely also suggests regions of disorder because the downward trend towards the tail of the data is not overly strong when compared to other folded RNA in solution^32, 37^ B2Δ81-124 does not show as strong of a trend, which reinforces the previous SHAPE data suggesting that this RNA species is not strongly folded and tends towards having unstructured region(s) (Figure 2E). Finally, the data is represented as paired distance distribution functions, allowing for rough shape inference and measuring the maximum particle dimension (D_max_). Three RNA species, B2, B2J, and B2Δ96-105, have a very similar Dmax of ∼175 Å, consistent with the Rg measurements. B2Δ81-124 shows a much larger Dmax of 220 Å, indicating that it is likely oligomerized and might not accurately reflect a monomer.

We have represented *ab initio* reconstructions of each RNA in Figure 3(E-H). B2 and B2J are consistent in their three-dimensional reconstruction, displaying a similar tertiary architecture (Figure 3E and F). Both RNA species have a singular elongated portion that branches/fans out towards the right side (based on the figure). Interestingly, while the dimensions (R_g_, D_max_) of B2Δ95-105 are remarkably similar to B2 and B2J, the overall three-dimensional reconstruction looks quite different. The branching/fanning of the RNA is no longer apparent, and the bends in the overall structure are far more evident (see Figure 3G). This change suggests that the small deletion does not significantly affect the physical measurements; however, the tertiary architecture has changed quite dramatically. Furthermore, the three-dimensional representation of B2Δ81-124 is significantly different compared to the other three RNA species and shares virtually no structural similarities (see Figure 3H and Table 1). The B2Δ81-124 tertiary conformation also does not appear to have any structural symmetry, suggesting that the dimerized RNA might fold into a brand-new conformation and not simply be the association between two structurally identical monomeric units.

### High-Resolution Computational Prediction of B2 Sine RNA Structure

We sought to understand better the three-dimensional organization of B2 RNA by coupling our SAXS experimental low-resolution envelopes with structure prediction computational pipeline. We, therefore, implemented a fragment-based Monte-Carlo simulation approach using the RNA modeling program ERNWIN^38^ to produce all-atom structures for B2 and its mutant variants. To model B2 RNA and its variants, we implemented experimentally determined secondary structure(s) and experimental one-dimensional SAXS data as co-restraints to generate an ensemble of all-atom B2 RNA structures that we found to be highly consistent with the SAXS data.

We modeled B2 WT RNA and generated an ensemble of atomistic structures that fit the pair-distance distribution function (P(r)) and the raw scattering data with a 1D correlation coefficient (*χ*_1*D*_) range of 1.7 to 2.7 (see Supplementary Figure 3A). By superimposing the generated models onto the SAXS-derived solution structure, we found that the computational model partially or fully matches the solution structure. To quantify the correlation between the 3D structure, we calculated the 3D structure correlation coefficient (SCC_3*D*_). We implemented a structure-based coarse-graining method to further improve the SCC and performed molecular dynamics (MD) simulations by considering SAXS density as a restraint motif. The SCC significantly increased while maintaining the *χ*_1*D*_ values. The best models show SCC_3*D*_ lies in the range of 0.70 to 0.75, showing a higher correlation with the SAXS-derived low-resolution solution structures (see Table 1). This close agreement between the one and three-dimensional SAXS data displays the method’s capability. The top 3 representative models of B2 RNA superimposed on SAXS-derived solution structure are presented in Figure 4A and rotated for clarity. Of all the best representative models, the most considerable variation in the all-atom structures lies within helices H4, H5, and H6 (Figure 4D). Model 1 shows that H5 and H6 can exist tightly packed together, while models 2 and 3 show H5 and H6 considerably more spread out, while H4 is relatively static in its position and orientation along with Domain I (nt# 1-72). Previous experiments show that nucleotides C76, C103, and U110 are involved in tertiary interactions through RNase V1 assays^24^. Interestingly, we do not see any tertiary interactions within our models of B2 WT RNA at positions C76, C103, and U110.

**Figure 4:**
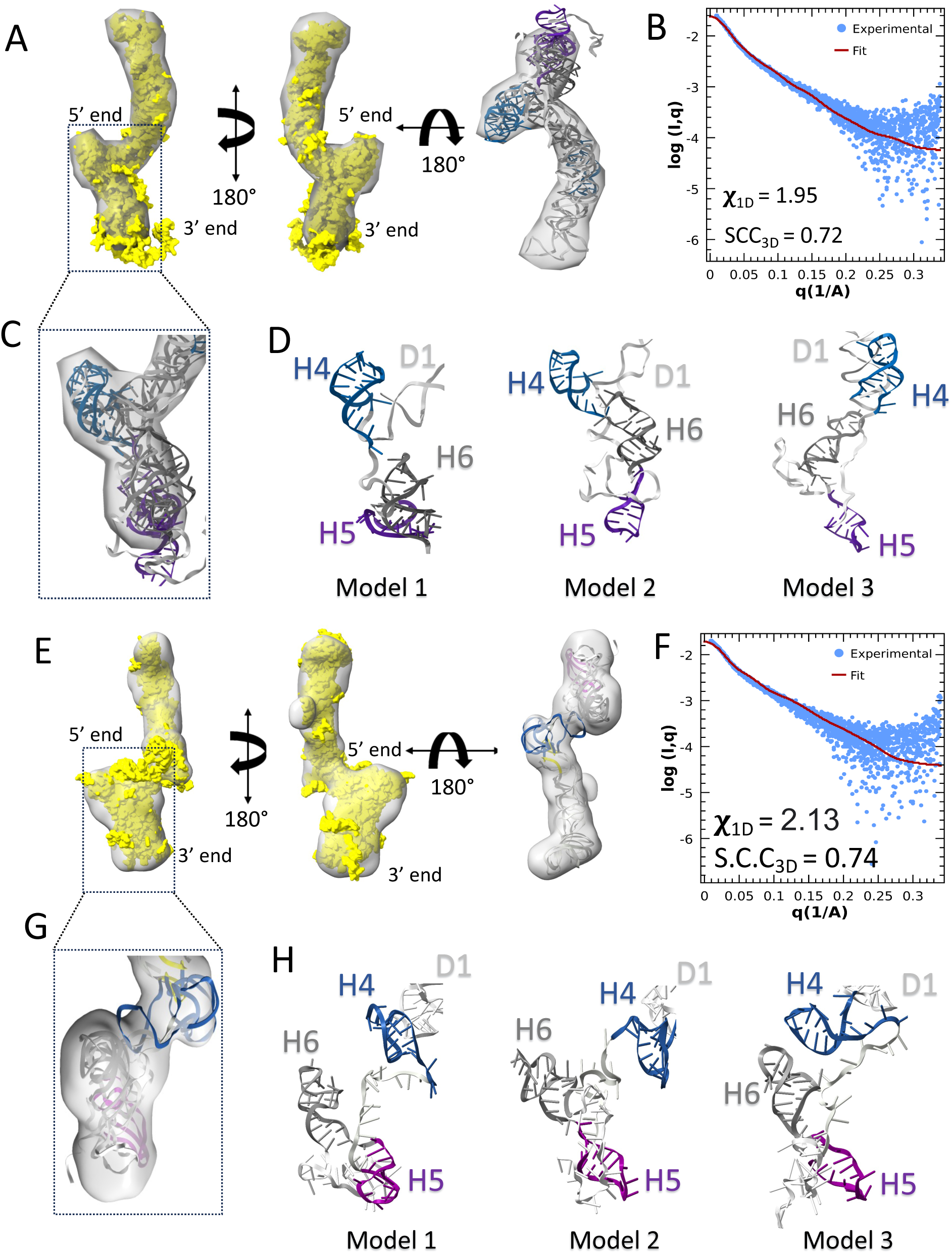
(A) A comparison of the averaged SAXS solution structure of full-length B2 (meshed black) and a computationally derived atomistic model (yellow) shows agreement. (B) The scattering intensity (Log I(q)) vs. q displays agreement between experimental SAXS data (blue) and simulated data calculated from the all-atom model (red) for B2 RNA. (C) Superposition of the top 3 model zoom-ins of highly dynamic loops in B2 RNA. (D) Three different configurations of loops, H4 (blue), H5 (purple), and H6 (gray), are shown for the top 3 models. (E) A comparison of the averaged SAXS solution structure of full-length B2J (meshed black) and a computationally derived atomistic model (yellow) shows agreement. (F) The scattering intensity (Log I(q)) vs. q displays agreement between experimental SAXS data (blue) and simulated data calculated using the all-atom model (red) for B2J RNA. (G) Superposition of the top 3 model zoom-ins of highly dynamic loops in B2J RNA. (H) Three different configurations of the dynamic loops of B2J RNA, H4 (blue), H5 (purple), and H6 (gray), are shown for the top 3 models.

We followed the same protocol to model B2 RNA variants and generated an ensemble of models for B2J, B2Δ96-105, and B2Δ81-124. As mentioned, B2J displays a similar tertiary structure to B2 RNA, which was also consistent in the all-atom models (see Figure 4E and Supplementary Figure 3B). We also observed a strong correlation between the raw scattering data and simulated SAXS data calculated from all-atom models, as shown in Figure 4F. Comparatively, we see conformational changes in the structure of helices H4, H5, and H6, the three most dynamic regions within B2 WT RNA. B2J shows a degree of variation with helix H4 (Figure 4H), where the H4 is very dynamic, whereas in B2 WT, H4 was relatively static in position and orientation. Alternatively, both H5 and H6 in B2J are reasonably consistent in orientation and position between the top 3 models (Figure 4H). This restructuring suggests that while B2J contains only two-point mutations compared to WT, these point mutations can cause conformational changes in the tertiary architecture, shifting the relative dynamics from flexible H5 and H6 helices in B2 cause helix H4 to move from a static state within B2 WT to a dynamic structure highlighted in Figure 4H. Therefore, we postulate that the shifting of tertiary architectural dynamics within helices H4, H5, and H6 caused by the two-point mutations at positions nt# 55 and 114 changes self-cleavage between B2J and B2 WT. As mentioned, nucleotides 45-55 are suggested to influence cleaving but show little difference in secondary structure analysis. However, comparing tertiary structure(s) of the same region, between B2 and B2J, we see a difference in a few representative models of B2J, whereas region nt 45-55 exists as a protruded loop compared. These regions could, therefore, be important for either protein binding or directly in self-cleavage.

The B2 deletion variant, B2Δ96-105, was observed to have a very different tertiary structure relative to the B2 (WT) and B2J RNA via SAXS experiments (Figure 2C and Figure 3G), and these changes are also evident in the all-atom models. The SCC_3*D*_ between the predicted structures and experimental SAXS data is 0.72 (Figure 5B), suggesting a strong correlation and high confidence in the models. Overall, we observe a single major conformation for B2Δ96-105 due to its highly base-paired secondary structure, further suggesting the deletion reduces the conformational space of B2Δ96-105 (Figure 5A) compared to WT B2 and B2J. Although we observed significant consistency between the models, we could not fit the disordered region at the tail 3’ end of B2Δ96-105 within the experimental SAXS envelope, suggesting extreme flexibility of this region. We, therefore, purposefully excluded this region from the models (see Figure 4 C).

**Figure 5:**
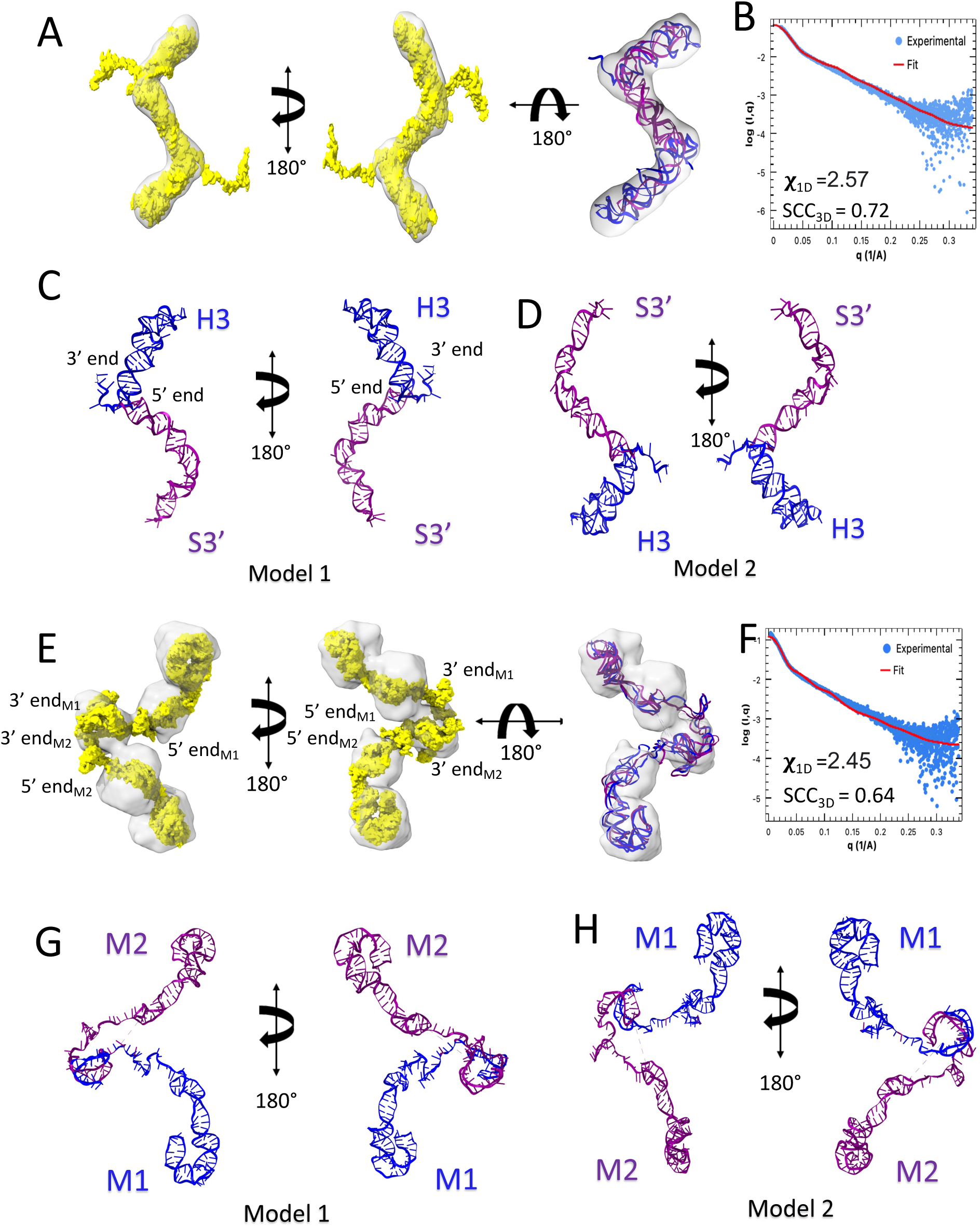
(A) A comparison between the averaged SAXS solution structure of full-length B2Δ96-105 (black) and a computationally derived atomistic model (yellow) shows agreement. Atoms outside the SAXS-derived solution structure indicate flexible regions. (B) The scattering intensity (Log I(q)) vs. q displays agreement between experimental SAXS data (blue) and simulated data calculated from the all-atom model (red) for B2Δ96-105 RNA. (C and D) Top models show a single major B2Δ96-105 conformation, likely due to its highly base-paired secondary structure. This single major conformation suggests that the deletion reduces the conformational space of B2Δ96-105. The two arms, H3 (blue) and S3’ (purple), depict the symmetric fold of B2Δ96-105. (E) A comparison between the averaged SAXS solution structure of full-length B2Δ81-124 (black) and a computationally derived all-atom model (yellow) shows agreement. (F) The scattering intensity (Log I(q)) vs. angle displays agreement between experimental SAXS data (blue) and simulated data calculated from the all-atom model (red) for B2Δ81-124 RNA. (G and H) The top two all-atom models show two monomers, M1 (blue) and M2 (purple), demonstrating oligomerization of the RNA, leading to the increase in the size of the SAXS envelope.

Similar to the SAXS-derived conformation, we observe disordered all-atom models for B2Δ81-124. Initially, we assumed this to be caused by the significant lack of structure of the RNA and generated all-atom structures that, while unstructured, could still fit the data, but it resulted in much higher *χ*_1*D*_ and observed very poor correlation between the all-atom structure and SAXS-derived solution structure (see Supplementary Figure S4). We then hypothesized that oligomerization of the RNA could lead to the increase in the size of the SAXS envelope, so we shifted towards creating all-atom structures of a dimerization of B2Δ81-124. Surprisingly, we obtained a reasonable SCC_3*D*_ correlation score of 0.64 between our predicted structure(s) and the experimental SAXS-derived envelopes (Figure 5F). We attribute the dimerization to the extensive unstructured region of the B2Δ81-124 5’ end, which is considerably longer than the other RNA variants, even B2Δ96-105. Importantly, each of these B2 variants shows reduced self-cleavage compared to B2 WT, and each displays a difference in H4-H6 architecture. This information leads us to suggest there is a link between B2 ribozyme activity and the structural dynamics within these helical regions.

## Conclusion

Since the discovery of ribozymes, researchers have validated ∼15 different classes. They perform many functions, such as polymerizing amino acids to proteins, cleaving tRNA in the case of RNase P, and essential splicing reactions like the Groups I and II introns. Out of all the known ribozymes, only six are self-cleaving in nature. Recently, RNAs have piqued interest as base materials for several biomedical applications since RNAs’ unique structural features facilitate targeting the ribozymes’ catalytic activity and exploiting their catalytic functions to target other cellular RNAs. However, RNA structure information is needed to develop the RNA therapeutic potential as an appropriate function for any potential RNA motif. B2 has been classified previously as an epigenetic ribozyme^25^. Cellular stress modulates self-cleavage and interaction(s) with a known chromatin-modifying methyltransferase that targets lysine 27 of histone H3, EZH2. However, the complex they form upon cellular stress is independent of EZH2 catalytic activity, and therefore, it mostly leads to B2 cleavage in loci that need to be derepressed under stress. Similar to how group I and group II introns function, it is theorized that upon interacting with protein partners, catalytically competent B2 conformations are stabilized, promoting self-cleavage. Therefore, it is crucial to understand both the secondary and tertiary architecture of B2 in the context of being an epigenetic ribozyme. Here, we have resolved the B2 SINE ribozyme and its variant solution structure by combining experimental and computational techniques, including (i) transcription of B2 and its mutant, (ii) resolving secondary structure using chemical SHAPE probing, (iii) solution structure determination using SAXS, and (iv) constructed an ensemble of simulated solution structures based on experimental data. Our study resolves the solution structural ensemble of B2 SINE and elucidates the effect of point mutations and deletions of regions of the B2 RNA on its secondary and tertiary architecture. We find that the removal of the main cleavage site rearranges the secondary structure and leads to a reduction in conformational space. Removing larger regions surrounding this site leads to oligomerization. Two point mutations, as seen in B2J, lead to more subtle changes, in the tertiary architecture of B2 WT, shifting the relatively static state of transcriptionally active loops to a more dynamic state.

Regarding point mutations, our SHAPE analysis found similar secondary structures for the wild-type B2 RNA and B2J RNA, which include two-point mutations at A55C and U114C. Our SAXS and all-atom solution structures validate this and, in addition, suggest single point mutations induce a local change in the RNA tertiary structure, where domain H4 becomes more disordered with the mutation. Because the SHAPE reactivity profiles are similar for wild type and mutant, whereas the SAXS and computational modelling analyses show significant differences, we believe the secondary structure is unchanged, while some tertiary contacts are remodeled. Reports show that the minimal span of positions 81-131 in B2 RNA can inhibit transcription by more than 50%^24^. Our results suggest that this transcription inhibition could be due to the changes in the tertiary structure and mobility, but we acknowledge that additional studies will be required to support this hypothesis. In addition to point mutations, we have also studied deletions of larger sections of the B2 RNA. Deletion of a short, 10-nucleotide region of RNA (positions 96-105) causes a rearrangement of the secondary structure and tertiary structure. The SAXS and computational modelling analysis of this deletion suggests reduced conformational flexibility. A longer, 44-nucleotide deletion (positions 81-124) causes a loss of secondary structure and surprisingly results in the oligomerization of the RNA monomer. Lastly, our protocol predicts the 3D structural ensemble of B2 RNA with a high correlation between our ensemble of all-atom models and our experimental SAXS densities. This structural study of B2 and its variants will provide information for future studies, such as the association of B2 RNA with known interacting proteins like EZH2^39^ and the change in dynamic associations of RNA-protein complexes based on ncRNA active sites. By integrating biochemistry, structural biology, and computational methods, we unravel the in-solution conformations of the B2 RNA and open the possibility for further studies of other ncRNA and mRNA systems.

## Material and methods

### Transcription and Purification of B2 Sine RNA

Linear template DNAs containing a T7 promoter and B2 different sequences used in this study were obtained from a linear gBlock (IDT) containing the following B2 WT sequence (5’ to 3’). The gBlock was subjected to polymerase chain reaction using Ex Taq high fidelity DNA Polymerase (Takara Bio USA, # RR001A) and the following primers 5’-TAATACGACTCACT ATAG-3 and as a reverse primer for the B2 RNA PCR template, the following sequence 5’-TTTTTTTTTA AAGATTTTTTATTTATTATATGTAAGTACA-3’. *In vitro* transcription reactions were run using the AmpliScribe T7 High Yield Transcription Kit (Lucigen AS3107) for 4 hours at 42°C, followed by a 15 min incubation with DNase I at 37°C. Reaction products were diluted to 1 mL with size exclusion buffer (8 mM MOPS pH 6.5, 100 mM KCl, 0.1 mM EDTA) and purified in an AKTA Pure 25 system on a Superdex 200 Increase 10/300 GL column (GE Healthcare). Fraction aliquots were resolved in 8M Urea TBE polyacrylamide gel electrophoresis, and their respective eluates were pooled if they contained the desired full-length product.

### B2 ribozyme activity assays

Activity assays were performed by pre-folding B2 RNA in ribozyme buffer (5 nM Tris pH 7.9, 300 mM NaCl, 0.5 mM MgCl_2_, 0.02 mM EDTA, 0.01% NP40, 1% Glycerol, and 0.2 mM DTT). RNAs were heated for 1 min at 55°C and cooled to 37°C at a rate of 0.1 °C/s. After folding, 200 nM B2 was subjected to activity assays at 37°C in ribozyme buffer. Reactions were inactivated at the indicated times with 50% Formamide, and the reaction products were separated and analyzed by 8M Urea TBE polyacrylamide electrophoresis at 45°C and a constant 30 W.

### Design of RNA for chemical probing experiments

The synthetic DNA template corresponding to the B2 sine RNA transcript (AC020972.3 121488-121665) was placed in the center of an RNA “structure cassette” (25) to facilitate analysis of 2’-O-adduct formation by the chemical probing reaction. Each helix in the cassette contains a stable UUCG tetraloop (26) to enforce the designed fold (of the cassette) and eliminate interference with the RNA sequence of interest. The sequences of 5’ and 3’ structure cassettes are provided below in Table 2.

**Table 2:**
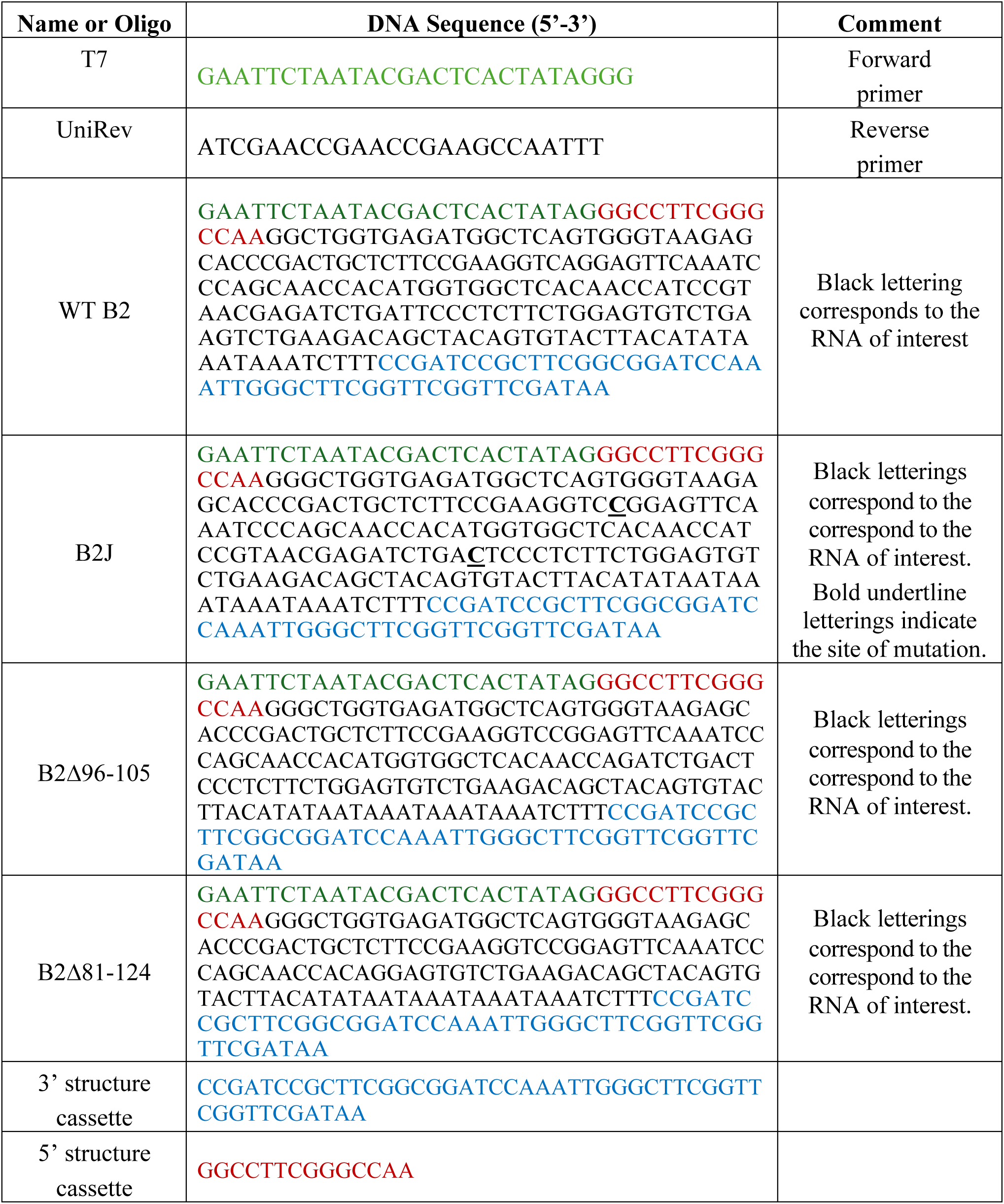
List of oligos and DNA constructs for chemical probing

### *In vitro* transcription of B2 and B2J RNA for chemical probing

We purchased double-stranded DNA templates (with 5’ and 3’ structure cassettes) for *in vitro* transcription from Integrated DNA Technologies (Coralville, IA). The desired RNA is transcribed via run-off transcription utilizing the AmpliScribe T7 High Yield Transcription Kit (Biosearch Technologies, Catalog # AS3107). The resulting RNA is precipitated by adding a mixture of 3 volumes of anhydrous ethanol and 1 volume of 7M ammonium acetate and recovered through centrifugation. The RNA pellet is then resuspended in nuclease-free water and purified by RNAClean XP bead (Beckman Coulter, Catalog # A63987) by manufacturer protocol. RNA is produced fresh (not older than 6 hr) to minimize the chance of self-cleaving. The integrity of the prepared RNA is validated by dideoxy sequencing before probing experiments.

### SHAPE probing

Thirty picomoles of the transcribed RNA were heat-denatured at 95°C and flash-cooled on ice. The refolding of RNA was performed in a buffered solution containing 50 mM HEPES (pH 8.0), 100 mM KCl, and 0.5 mM MgCl2 at 25°C. The reagent 1-methyl-7-nitroisatoic anhydride (1M7) was synthesized in-house as described by Mortimer et al.^40^. A fresh 1M7 SHAPE reagent solution in DMSO (at a final concentration of 6 mM) is added to the refolded RNA and incubated at 25°C for 10 min. A blank set is also performed in parallel by adding an equal volume of neat DMSO to the refolded RNA and incubating it similarly. Post incubation, the modified RNA is recovered via ethanol precipitation.

### Primer extension

The modified RNA is resuspended in 7.75 *µ*L of nuclease-free water and incubated successively at 65°C for 1 minute, 45°C for 5 minutes, and in ice for 1 min with 1 *µ*L of 1.5 *µ*M (1.5 pmole) of 5’-Alexa-488 labeled Unirev primer (Integrated DNA Technologies (Coralville, IA)). The primer extension mix contained 500 *µ*M of each dNTP, 10 mM DTT, 1X SSIII FS buffer, and 200 U/ *µ*L of SuperScript™ III Reverse Transcriptase (Invitrogen). A 15 *µ*L primer extension reaction is initiated by incubating the mixture at 55°C for 1.25 hours. Similarly, A and G sequencing reactions were performed on thermally denatured unmodified RNA. The primer extension mix was supplemented with 500 *µ*M ddTTP and ddCTP (Cytiva) for A and G sequencing reactions. Primer extension reactions were desalted by spinning in the P-6 micro-biospin column (BioRad). Each sample was diluted in deionized formamide (Hi-Di™ Formamide, Thermo Fischer Catalog # 4311320) in a 1:20 ratio (by volume) and heated to 95°C for 3-4 minutes. The samples were electrokinetically injected (30 s at 6 kV) onto an ABI Prism 3100 Avant quad-capillary instrument, and a fluorescence electropherogram was collected at 14 kV.

### SHAPE Data processing

Each electropherogram is processed as per the protocol reported earlier^41^. Briefly, each data set (SHAPE and sequencing) is aligned and integrated using in-house Perl scripts to simultaneously fit multiple Gaussian peaks to the traces.

### Small-angle X-ray scattering analysis

Sample data were collected at Diamond Light Source Ltd. synchrotron (Didcot, Oxfordshire, UK) utilizing the B21 SAXS beamline. An Agilent 1200 HPLC system (Agilent Technologies, Stockport, UK) system attached upstream to ensure sample monodispersity^42^ was used in conjunction with a Shodex KW403-4F (Showa Denko America Inc., New York, NY, USA) size exclusion chromatography column and a specially produced flow cell. RNA samples were injected into the equipment in 1X B2 Buffer (5 mM Tris-HCl [pH 7.9], 10 mM NaCl, 0.01% NP-40, 0.02 mM EDTA, 0.2 mM DTT, 0.5 mM MgCl_2_ and 1.5% glycerol) with a constant flow rate of 0.160 mL/minute. Eluted RNA samples were exposed to X-rays with three 3-second exposure times and ∼620 frames. SAXS data for B2, B2J, and B2Δ96-105 and B2Δ81-124 RNA samples were collected at 1.6 mg/mL for B2 and 1.5 mg/mL for the other three RNA.

Scattering data analysis was performed using the ATSAS suite 3.0^43^. Firstly, Chromixs was utilized to subtract the buffer contribution from each RNA sample peak^44^. Next, Guinier analyses (q^2^ vs. ln(I(q))) were performed on each RNA data set to obtain the reciprocal radius of gyration (R*_g_*) while additionally inferring sample quality^45^. The general folding of RNA molecules was evaluated using a dimensionless Kratky analysis (qR*_g_* vs. qR*_g_*^2^*I(q)/I(0))^46^ Using paired-distance distribution P(r) analysis via GNOM, real space Rg and particle dimension maxima (Dmax) were calculated for each RNA sample^47, 48^. 20 models for all B2 WT and mutant constructs, were generated from P(r) derived information and using DAMMIN^49^. Finally, utilizing DAMAVER, the models were averaged and filtered via DAMFILT, producing a representative model^49, 50^.

### Atomistic structure modeling

To model atomistic RNA structure, we performed coarse-grained RNA fragment simulation based on the Monte Carlo algorithm to build the tertiary structure. A total of 20,000 simulation steps were performed to get best fitting with the experimental SAXS data. Coarse-grained RNAs were converted to all-atom structures at every 1000 steps. Each all-atom structure was then used to calculate the correlation between the pair-distance distribution function of the proposed computational tertiary structure and the experimental SAXS distance distribution. After completing the sampling steps, we calculated theoretical SAXS data and compared it to the experimental SAXS one-dimensional data and filtered the top predicted structure with *χ* values in the range of 1-3. We further performed the restrained molecular dynamics simulation using MDfit^51^ to improve the structure correlation coefficient between the average solution structure generated by SAXS and the all-atom models by considering density as a restraint motif.

## Data Availabilty

SAXS data has been deposited to SASDB under accession number SASDV63, SASDV93, SASDV83, and SASDV73 for B2 WT, B2J, B2Δ96-105, and B2Δ81-124 respectively.

