## Supplemental Figure 1 for "Cleavage region organizes the structural architecture of the B2 SINE ribozyme"

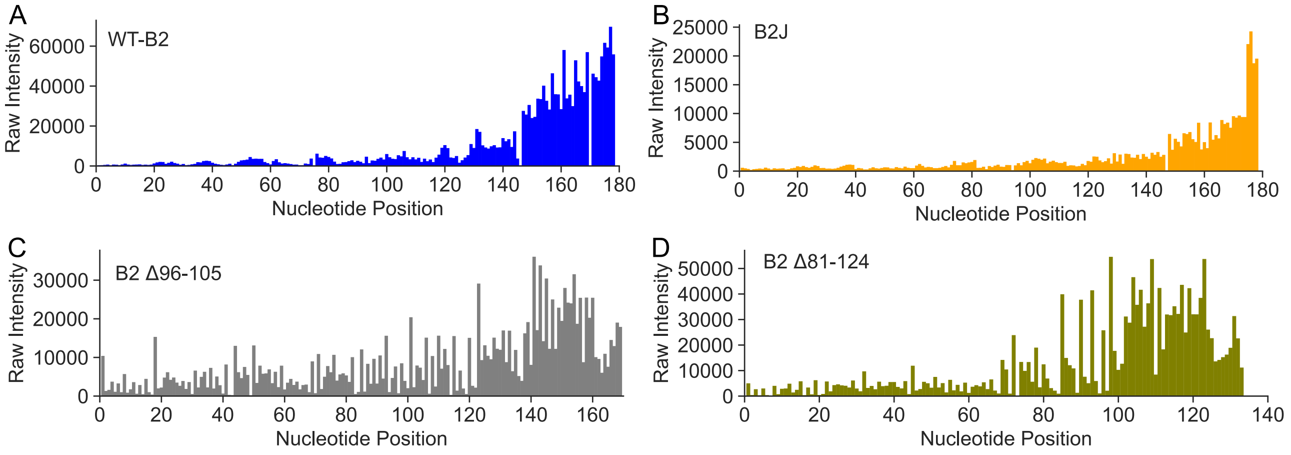


**Supplementary Figure S1**: Raw SHAPE electropherogram of (A)WT B2 RNA (B) B2J (C) B2Δ96-105 and (D) B2Δ81-124. The electropherogram corresponds to the SHAPE probing experiment of full-length RNA sequences for all the cases. The low SHAPE reactive regions indicate the presence of structured regions, whereas nucleotides with high SHAPE reactivities indicate interhelical junctions.

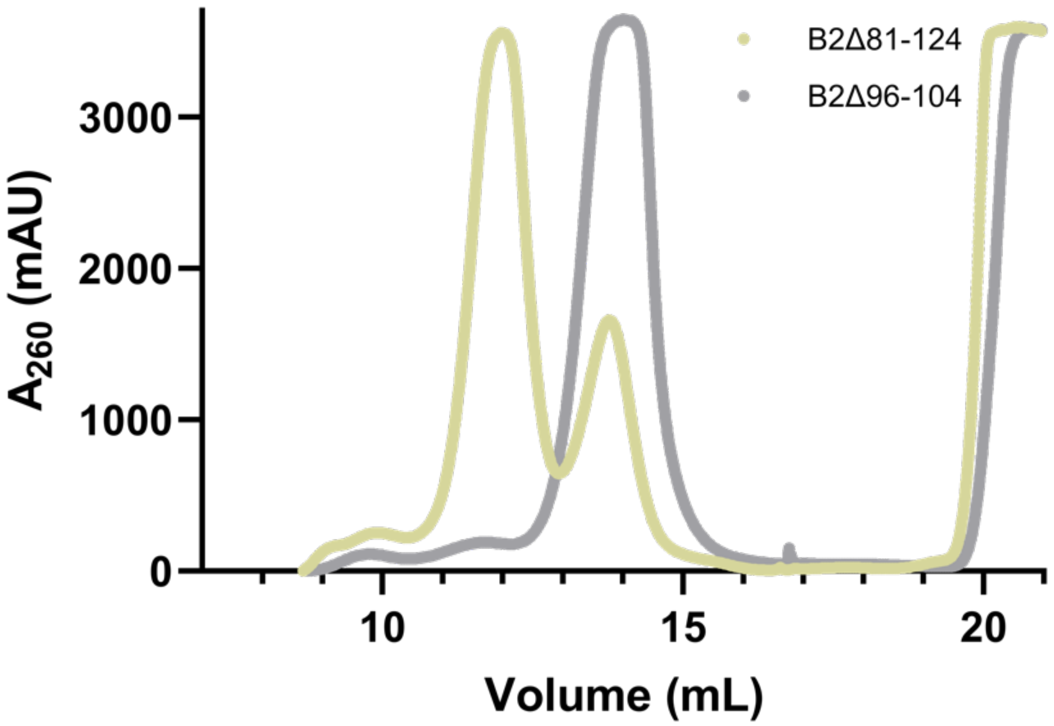


**Supplementary Figure S2**: Size exclusion chromatogram (Superdex 200 increase, 10/300 GL) showing the elution profiles for B2Δ96-105 and B2Δ81-124 purification which highlights the lack of any self-cleavage products that are prominent in the B2 wildtype purification (Figure 1B).

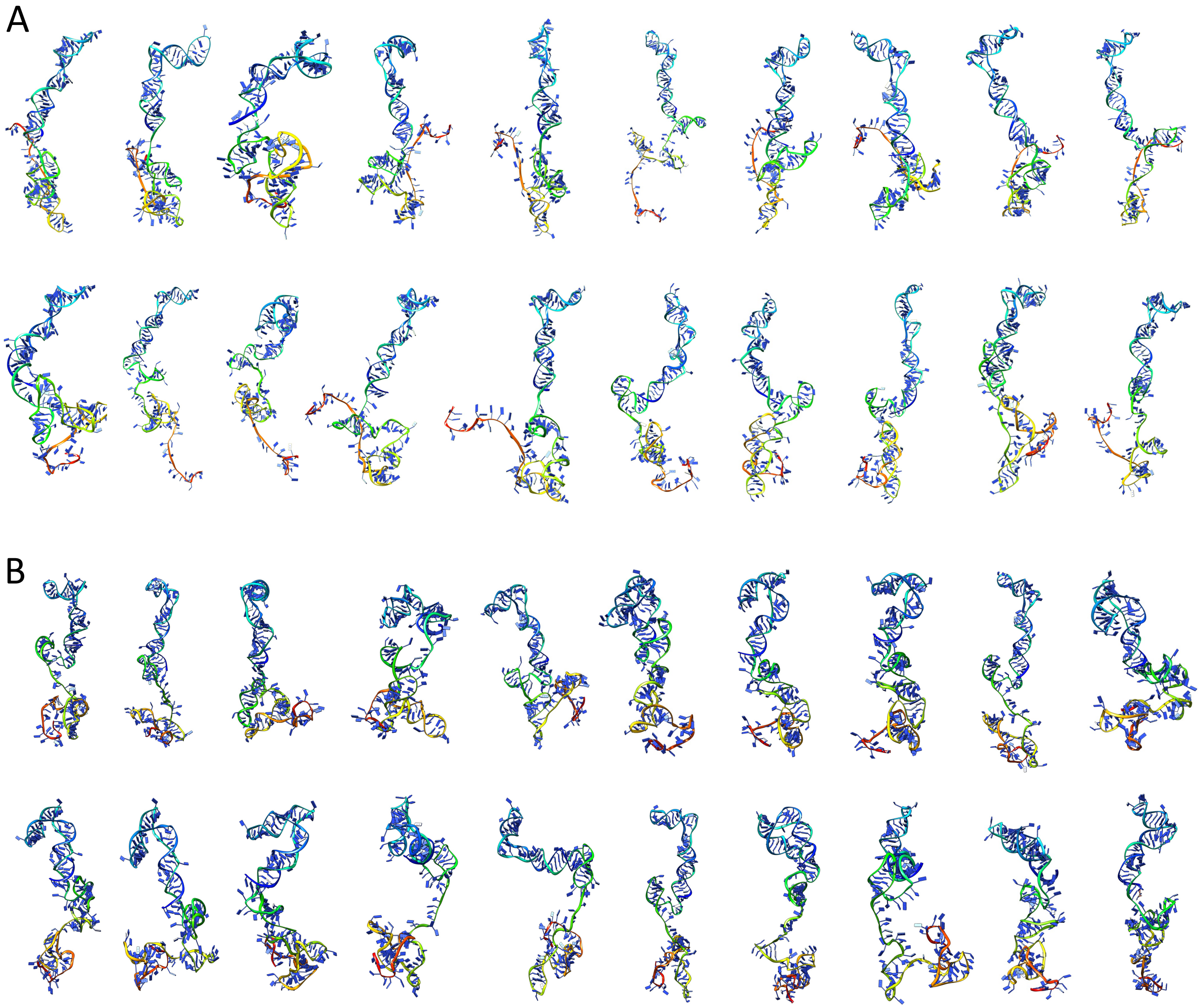


**Supplementary Figure S3**: 20 top-ranked atomistic models of full length at 0.5 mM $Mg^{2+}$ (A) B2 RNA and (B) B2J RNA. Each all-atom was generated at every 1000 steps using the RNA modeling program ERNWIN.

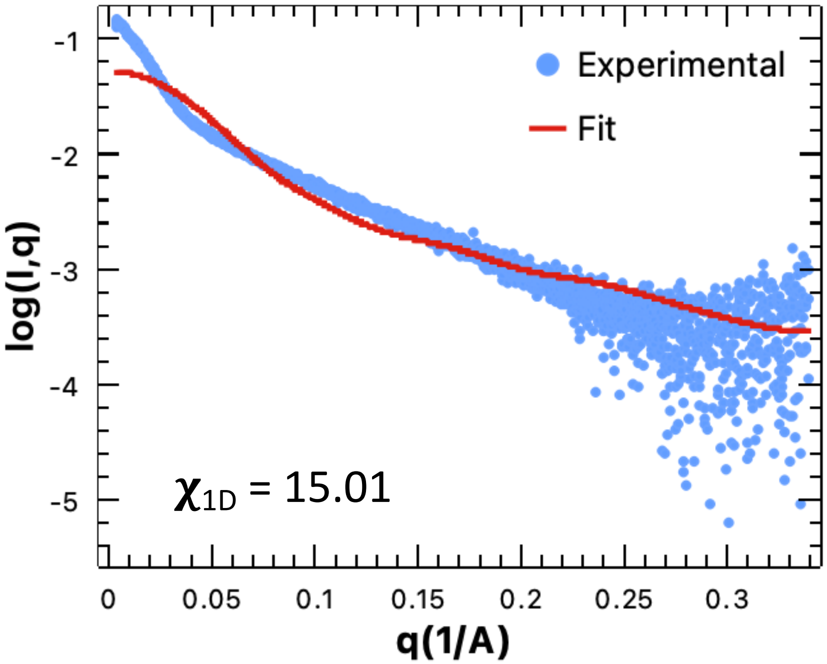


**Supplementary Figure S4**: The scattering intensity (Log I(q)) vs. q between experimental SAXS data (blue) and simulated data calculated from the all-atom model (red) for B2Δ81-124 RNA Monomer (M1). The all-atom monomer model resulted in much higher $\chi_{1D}$ between the all-atom structure and SAXS-derived solution structure.
